# Late-life Rapamycin Treatment Enhances Cardiomyocyte Relaxation Kinetics and Reduces Myocardial Stiffness

**DOI:** 10.1101/2023.06.12.544619

**Authors:** Akash D. Chakraborty, Kristi Kooiker, Kamil A. Kobak, Yuanhua Cheng, Chi Fung Lee, Maria Razumova, Henk Granzier H, Michael Regnier, Peter S. Rabinovitch, Farid Moussavi-Harami, Ying Ann Chiao

## Abstract

Diastolic dysfunction is a key feature of the aging heart. We have shown that late-life treatment with mTOR inhibitor, rapamycin, reverses age-related diastolic dysfunction in mice but the molecular mechanisms of the reversal remain unclear. To dissect the mechanisms by which rapamycin improves diastolic function in old mice, we examined the effects of rapamycin treatment at the levels of single cardiomyocyte, myofibril and multicellular cardiac muscle. Compared to young cardiomyocytes, isolated cardiomyocytes from old control mice exhibited prolonged time to 90% relaxation (RT_90_) and time to 90% Ca^2+^ transient decay (DT_90_), indicating slower relaxation kinetics and calcium reuptake with age. Late-life rapamycin treatment for 10 weeks completely normalized RT_90_ and partially normalized DT_90_, suggesting improved Ca^2+^ handling contributes partially to the rapamycin-induced improved cardiomyocyte relaxation. In addition, rapamycin treatment in old mice enhanced the kinetics of sarcomere shortening and Ca^2+^ transient increase in old control cardiomyocytes. Myofibrils from old rapamycin-treated mice displayed increased rate of the fast, exponential decay phase of relaxation compared to old controls. The improved myofibrillar kinetics were accompanied by an increase in MyBP-C phosphorylation at S282 following rapamycin treatment. We also showed that late-life rapamycin treatment normalized the age-related increase in passive stiffness of demembranated cardiac trabeculae through a mechanism independent of titin isoform shift. In summary, our results showed that rapamycin treatment normalizes the age-related impairments in cardiomyocyte relaxation, which works conjointly with reduced myocardial stiffness to reverse age-related diastolic dysfunction.

## Introduction

Aging is a major risk factor for developing heart failure (HF) and other cardiovascular diseases such as coronary artery disease and stroke^1^. Approximately 1% of individuals aged over 50 years are affected by HF, and this number doubles with each decade of life, making HF the major cause of mortality in the elderly^1^. This is a matter of increasing concern in the United States, where the population aged 65 and over increased from 40 million in 2007 to 51 million in 2017 and is projected to reach 95 million in 2060^2^. Cardiac aging in both humans and mice is characterized by a progressive decrease in diastolic function and an increase in left ventricular (LV) hypertrophy^3^. Diastolic dysfunction is also a characteristic of HF with preserved ejection fraction (HFpEF), a form of HF that is prevalent in the elderly. HFpEF is defined clinically by signs of HF combined with preserved left ventricular (LV) ejection fraction (EF) ^4,5^. HFpEF patients have reduced cardiac output associated with impaired diastolic LV filling, and this results in exercise intolerance and contributes to frailty^5^.

A leading target of interventions for slowing aging and improving healthspan is the small molecule, rapamycin, an FDA-approved drug that directly inhibits the mechanistic target of rapamycin (mTOR) Complex I (mTORC1)^6–8^. Inhibition of mTORC1 has wide-ranging effects *in vivo*, including altering protein synthesis, inhibiting cell growth, and stimulating stress response mechanisms and autophagy^9^. Rapamycin treatment extends lifespan and healthspan of C57BL/6 mice or genetically heterogeneous mice in a dose- and sex-dependent manner^7,8,10,11^. We and other have previously demonstrated the beneficial effects of rapamycin on cardiac aging^12–15^. In particular, 10-week rapamycin (at a 14-ppm dose) treatment starting at late-life reverses pre-existing age-dependent diastolic dysfunction and cardiac hypertrophy in old female mice^12,13^. The improved diastolic function is accompanied by reduced oxidative damage^13^, increased proteome half-lives^13^, and transient increases in autophagy and mitochondrial biogenesis^12^. In a later study, we showed that late-life rapamycin also improves diastolic function of male mice when given a 42-ppm dose^15^. However, how rapamycin treatment impacts the molecular and cellular mechanisms regulating diastolic function have not been established.

Diastolic function is controlled by active relaxation of cardiomyocytes and passive stiffness of the myocardium^16,17^. Cardiomyocyte relaxation is controlled by the interplay of two macromolecular systems: membrane bound Ca^2+^ handling proteins to send the signal to start and stop contraction, and sarcomeric proteins for force generation and contraction regulation by Ca^2+18,19^. Passive stiffness of the myocardium is controlled by mechanisms such as extracellular matrix remodeling, titin isoform shift and titin phosphorylation^16,17,20–24^. Quarles et al. showed that rapamycin reduces the age-related increase in passive stiffness of the myocardium^15^. The effects of rapamycin on active cardiomyocyte relaxation and the precise molecular mechanisms of rapamycin mediated reduction in passive myocardial stiffness remain unknown. Identifying the mechanisms by which rapamycin improves diastolic function in the aging heart will advance our understanding on its therapeutic potentials in cardiac aging and HFpEF.

In this study, we investigated the effects of late-life rapamycin treatment on different molecular and cellular mechanisms that regulate diastolic function. We demonstrated that 10-week rapamycin treatment rescues age-related impairments in cardiomyocyte relaxation and reduces myocardial stiffness to improve diastolic function in old murine hearts. We detected a reversal in age-dependent derangements in Ca^2+^ sensitivity, cardiac myofilament properties and relaxation kinetics by rapamycin treatment for 10 weeks. By dissecting the molecular mechanisms underlying rapamycin-mediated diastolic function improvement in the aging heart, we aim to shed light on a newer dimension of mechanisms for reversal of age-related diastolic dysfunction.

## Methods

### Animal handling and Rapamycin treatment

Young (3-5-month-old) and old (23-25-month-old) C57BL/6J female mice were obtained from the National Institute of Aging Aged Rodent Colonies. Female mice were used as rapamycin offers a greater extent of protective effects against aging in female mice ^7^ and previous studies demonstrated rapamycin-induced reversal of diastolic dysfunction in old female mice^12,13^. All mice were handled according to the guidelines of the Institutional Animal Care and Use Committee (IACUC) at the Oklahoma Medical Research Foundation and the University of Washington.

Young control mice were fed with regular chow diet (LabDiet PicoLab Rodent Diet 20, #5053). Old mice were randomly assigned to two groups and were fed with: 1) rapamycin diet containing EUDRAGIT encapsulated rapamycin (from Rapamycin Holdings, San Antonio TX) at 14 ppm or 2.24 mg/kg/day, or 2) control diet containing EUDRAGIT encapsulation alone for 10 weeks^15^. Custom diets with EUDRAGIT or EUDRAGIT encapsulated rapamycin were synthesized by LabDiet.

For multicellular and myofibril mechanical measurements, mice were euthanized by i.p. injection of a lethal dose (0.1 mL) of pentobarbital (Beuthanasia-D). The heart was immediately removed and processed for downstream analyses. For cardiomyocyte isolation and biochemical analyses, mice were anesthetized by isoflurane and heart was excised for downstream processing.

### Isolation of adult mouse cardiomyocytes

For cardiomyocyte isolation, mice were injected with heparin, anesthetized by isoflurane and rib cages were cut open and hearts were immediately excised and perfused with cardiomyocyte isolation buffer and cardiomyocytes were isolated as previously described^25^. The standard cardiomyocyte isolation buffer contained: 120 mM NaCl, 5.4 mM KCl, 0.33 mM Na_2_HPO_4_, 0.5 Mm MgSO_4_.7H2O, 30 mM Taurine, 10 mM BDM, 25 mM HEPES, 22 mM Glucose, pH 7.4 with NaOH. For myocyte isolation, collagenase (2.4 mg/ml, Worthington biochemicals) and Protease XIV (0.2 mg/ml, Qiagen) were used in the presence of fetal bovine serum.

### Cardiomyocyte contractility and calcium measurement

Isolated cardiomyocytes were briefly incubated with 0.2 µM Fura-2-acetoxymethyl ester (Fura-2AM, Invitrogen), a calcium sensitive, ratiometric fluorescent dye, for 20 minutes at room temperature and kept in dark. Cardiomyocytes were washed in standard isolation buffer for 20 minutes to allow dye de-esterification. An IonOptix system was used to conduct the experiments, including the Xenon arc lamp, hyperswitch and myopacer for cardiomyocyte stimulation and fluorescence excitation, myocam-S for the measurement of sarcomere length, and a fluorescence system interface to integrate the different components (IonOptix Ltd). The Xenon arc lamp and fluorescence hyperswitch containing a galvanized mirror were used to alternate between wavelengths of 340 and 380 nM at a high frequency. Ca^2+^ fluorescence was recorded at 510 nm. Ca^2+^ traces represent the ratio of calcium bound: calcium free Fura-2-AM dye, and hence changes in free intracellular Ca^2+^ concentration. Myocytes were paced at 1 Hz and measurements of sarcomere length were made in isolation buffer containing 1.2 mM CaCl_2_. Both sarcomere shortening and Fura-2 fluorescence measurements were made simultaneously from the same cardiomyocyte.

### Western blotting

Pulverized cardiac tissues were homogenized in RIPA buffer (Sigma) with protease and phosphatase inhibitor cocktail (Halt, ThermoFisher). Protein concentrations of samples were determined by BCA assay (ThermoFisher), and equal amounts of protein (15 µg per sample) were loaded for SDS-PAGE using the Criterion system. Proteins were transferred onto PVDF membrane using Criterion Blotter (Biorad). The total protein stain was visualized using Revert total protein stain. Blots were blocked in 5% BSA-TBST. Primary antibodies were diluted using 5% BSA-TBST. Antibodies from the following companies were used for Western blot analysis: RyR2 (1:1000, abcam), RyR2-pS2808 (Badrilla, 1:1000), SERCA2 (1:5000, Badrilla), CSQ2 (1:3000, Cell signaling technology), NCX1 (1:1000, Cell signaling technology), MyBP-C-pS282 (1:2000, ALX), MyBP-C (1:1000, Santa cruz), TnI-pS23/24 (1:1000, Cell signaling technology), Total TnI (1:1000, Cell signaling technology), PKCα-pS567 (1:200, Santa cruz), Total PKCα (1:200, Santa cruz). Antibodies were diluted in 5% BSA-TBST. Protein bands were visualized with chemiluminescence assay (Pierce) with secondary antibodies coupled with HRP using the G-Box imaging system. The protein abundance was analyzed by densitometry with ImageJ and Image studio lite tools.

### Demembranated multicellular mechanics

Experiments were performed as described in Moussavi-Harami *et al*^26^. In brief, excised hearts were demembranated overnight at 4 °C in 50:50 (vol:vol) glycerol:relaxing solution (in mM: 100 KCl, 10 imidazole, 2 EGTA, 5 MgCl_2_, and 4 ATP) containing 1X protease inhibitor cocktail (Sigma-Aldrich P8340) and 1% triton X-100. Hearts were transferred to fresh glycerol:relaxing solution without triton X-100, then stored at -20°C for up to one week. Left ventricular multicellular tissues were dissected out, secured between aluminum t-clips, and then mounted between a motor (Aurora Scientific, Model 312B) and force transducer (Aurora Scientific, Model 403A). Preparations were moved to a bath containing experimental relaxing solution (pCa 9.0) at pH 7.0 at 15 °C containing (in mM): 15 phosphocreatine, 15 EGTA, 80 MOPS, 1 free Mg^2+^, 10^-9^ Ca^2+^, 1 DTT, and 5 Mg_2_ATP. For calcium sensitivity curves, the same components were used as for the experimental relaxing solution, with varying levels of free calcium from pCa 4.0 to 6.2 (pCa = -log[Ca^2+^]). Preparations were lengthened to sarcomere length of ∼2.3 µm then moved between solution baths where force was measured. After measuring force at each pCa and then returned to pCa 9.0, the length was reset to just above slack (L_0_), then sequentially stretched at 4% increments up to 24% L_0_. We scaled all data to the average force of young mice at 24% length change for each experiment to compare across experimental groups.

### Myofibril mechanical measurement

Small myofibril bundles were prepared from demembranated left ventricular wall tissue as previously described^27^. Briefly, bundles were rinsed twice in Rigor solution containing 2 mM DTT and 1:200 dilution of protease inhibitor (Sigma-Aldrich, St. Louis, MO) before being homogenized for 1 or 2 pulses of 30s at high speed, stored at 4°C, and used for up to three days. Experiments were performed on a custom set up as previously described^28^. In brief, myofibrils were mounted between two needles; one acted as a cantilever force transducer and the other as an inflexible mount attached to a piezo-electric computer controlled motor. A duel diode system was used to measure needle displacement and developed force was measured based on this displacement and the known stiffness of the needle. Needle stiffness was 2–7 nN/μm for this study. Relaxing (pCa=9.0) and activating (pCa=5.6) solutions were delivered to the mounted myofibril using a double-barreled glass pipette. Activation and relaxation data were collected at 15°C and fit with either single-exponential curves, linear coefficients, or 50% times as previously described^29^.

### Titin isoform analysis

SDS–agarose electrophoresis for analysis of titin isoforms was performed as previously described^30^. Briefly, pulverized cardiac tissues were solubilize in 40 parts of 8M urea and 40 parts of glycerol and analyzed by vertical SDS–agarose (1% agarose gel) electrophoresis. The gels were then stained with Coomassie blue and scanned for densitometry analysis of titin isoforms.

### Statistical analysis

Comparisons involving two experimental groups were analyzed by unpaired 2-tailed t-tests. Comparisons involving three groups were analyzed by one-way ANOVA with Tukey as a post-hoc test. All analyses were performed using GraphPad Prism 7.0. All data are expressed as mean ± SEM, and a *p*<0.05 was considered significant.

## Results

### Age-related impairments in cardiomyocyte contraction-relaxation kinetics are rescued by late-life rapamycin treatment

To better understand the effects of rapamycin treatment on active cardiomyocyte relaxation, we assessed the contraction-relaxation kinetics of cardiomyocytes isolated from young and old control mice and old mice with 10-week rapamycin treatment. **Figure 1A** shows the superimposed traces of sarcomere contraction-relaxation kinetics of cardiomyocytes from young, old control and old rapamycin treated mice. There was no difference in resting (baseline) sarcomere length in cardiomyocytes isolated from the three groups of mice **(Figure 1B)**. For sarcomere relaxation, old control cardiomyocytes displayed prolonged time to 90% relaxation (RT_90_) when compared to young cardiomyocytes **(Figure 1C)**, while time to 50% relaxation (RT_50_) and time to 10% relaxation (RT_10_) did not change between the groups **(Figure 1D-E)**. This age-related decline in cardiomyocyte relaxation kinetics (prolonged RT_90_) was completely rescued by rapamycin treatment **(Figure 1C)**. For sarcomere contraction, an age-related reduction in sarcomere fractional shortening was detected in old control cardiomyocytes and was normalized by 10-week rapamycin treatment to the levels of young cardiomyocytes **(Figure 1F)**. No differences in time to peak shortening (TPS) were observed in the three groups **(Figure 1G)**, but the time to 90% peak shortening (TPS_90_), 50% peak shortening (TPS_50_) and time to 10% peak shortening (TPS_10_) were significantly prolonged in old control cardiomyocytes when compared to young cardiomyocytes **(Figure 1H-J)**. These age-related impairments in contraction kinetics were partially normalized in old rapamycin-treated cardiomyocytes and there were no differences in TPS_90_, TPS_50 and_ TPS_10_ between old rapamycin treated cardiomyocytes and young cardiomyocytes **(Figure 1H-J)**.

**Figure 1:**
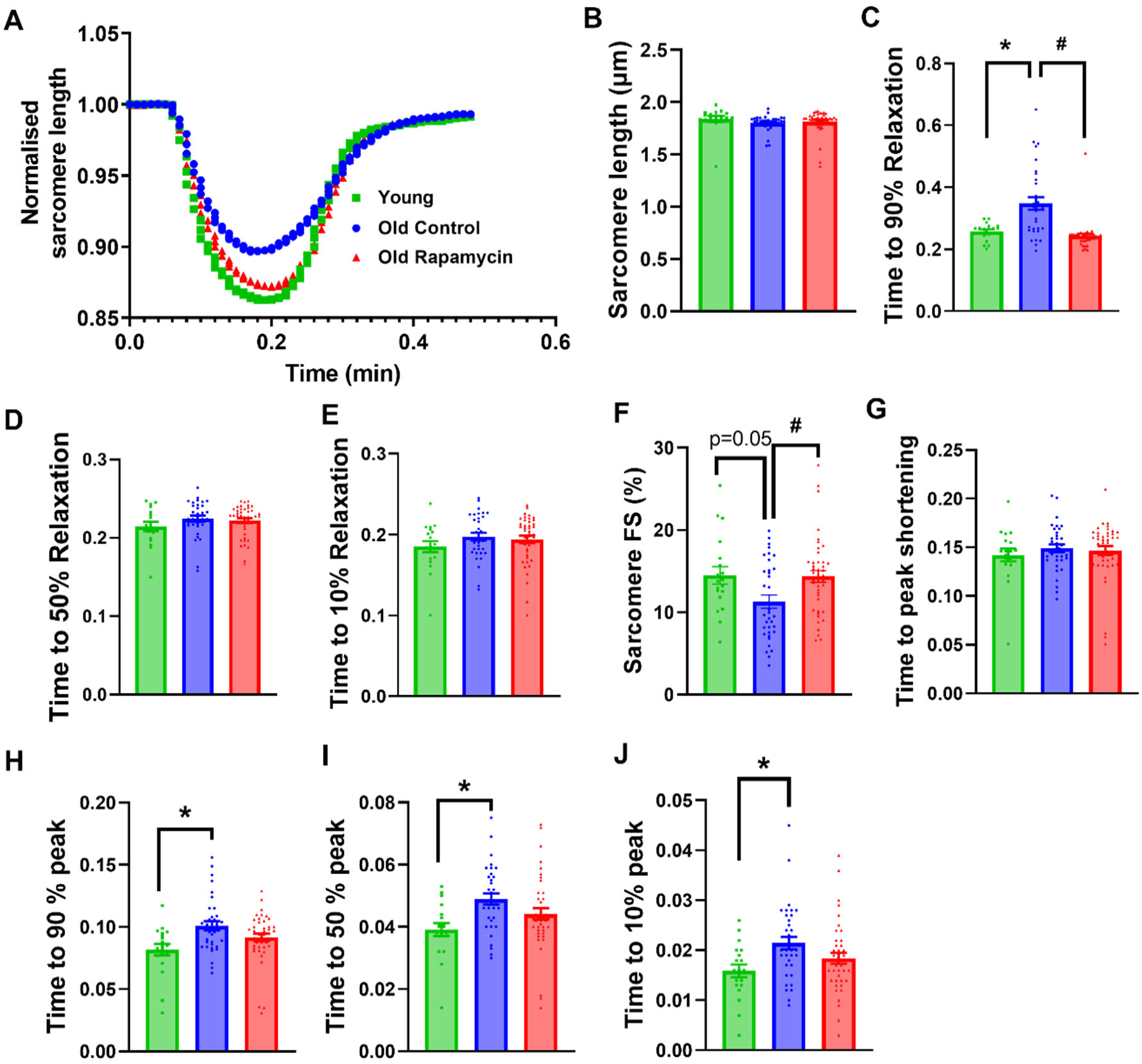
Rapamycin attenuates age-related impairments in cardiomyocyte contraction and relaxation kinetics. (A) Superimposed traces of averaged sarcomere contraction-relaxation kinetics of cardiomyocytes isolated from young, old control and old rapamycin treated mice. (B) Sarcomere length, (C) time to 90% relaxation, (D) time to 50% relaxation, (E) time to 10% relaxation, (F) sarcomere fraction shortening (FS), (G) time to peak shortening, (H) time to 90% peak shortening, (I) time to 50% peak shortening, and (J) time to 10% peak shortening of cardiomyocytes isolated from young, old control and old rapamycin treated mice. Data represented as mean±SEM; n=4-8 mice/group (number of cells measured – 19 cells for Young, 35 cells for Old Control and 43 cells for Old Rapamycin groups). *p<0.05 vs Young, #p<0.05 vs Old Control.

### Age-related dysregulation of Ca^2+^ handling is partially restored by rapamycin

In parallel to cardiomyocyte contraction-relaxation, we simultaneously measured intracellular Ca^2+^ transients in isolated cardiomyocytes to determine the contribution of calcium handling to the alterations in contraction-relaxation kinetics **(Figure 2A)**. Time to 90% Ca^2+^ decay (DT_90_) was significantly prolonged in old control and old rapamycin-treated mice compared to young controls, and DT_90_ trended to be lower in old rapamycin-treated mice compared to old controls **(Figure 2B)**. We detected no difference in time to 50% Ca^2+^ decay (DT_50_) between cardiomyocytes from young, old control and old rapamycin-treated mice (**Figure 2C**) and a trend toward longer 10% Ca^2+^ decay (DT_10_) in old control cardiomyocytes compared to young cardiomyocytes **(Figure 2D)**. The time to peak (TTP), time to 90% peak (TTP_90_), and time to 50% peak (TTP_50_) of Ca^2+^ transient was significantly prolonged in cardiomyocytes from old controls, but not in old rapamycin-treated mice, when compared to young controls (**Figure 2E-G**). TTP_90_ trended to be lower in old rapamycin-treated mice compared to old controls (**Figure 2H**). We detected no difference in time to 10% peak (TTP_10_) and Ca^2+^ transient amplitude between group (**Figure 2H and I**). These results suggest that impairments in the release and reuptake of cytosolic Ca^2+^ may contribute to the slower contraction-relaxation kinetics in old cardiomyocytes and these age-related impairments in Ca^2+^ handling are partially rescued by rapamycin treatment.

**Figure 2:**
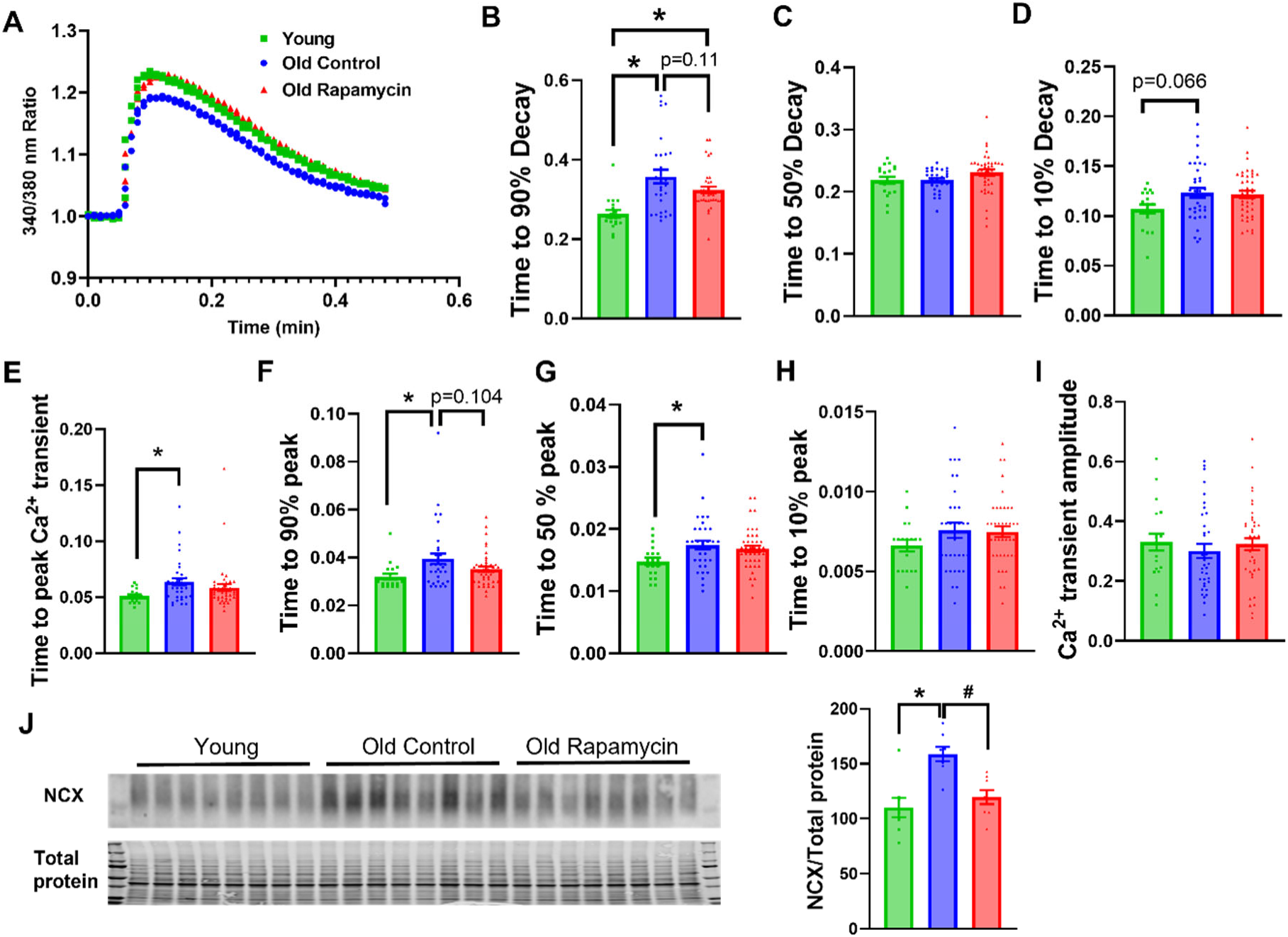
Aging results in prolonged cardiomyocyte Ca^2+^ transients and the derangements are partially normalized by rapamycin treatment. (A) Superimposed traces of averaged Ca^2+^ transients of cardiomyocytes isolated from young, old control and old rapamycin treated mice. (B) Time to 90% Ca^2+^ decay, (C) time to 50% Ca^2+^ decay, (D) time to 10% Ca^2+^ decay, (E) time to peak Ca^2+^ transient, (F) time to 90% peak Ca^2+^ transient, (G) time to 50% peak Ca^2+^ transient, (H) time to 10% peak Ca^2+^ transient, and (I) Ca^2+^ transient amplitude of cardiomyocytes isolated from young, old control and old rapamycin treated mice. Data represented as mean±SEM; n=4-8 mice/group (number of cells measured – 19 cells for Young, 35 cells for Old Control and 43 cells for Old Rapamycin groups). (J) Images of NCX western blot and total protein staining in heart tissues of young, old control and old rapamycin treated mice and the corresponding quantification of normalized NCX levels. Data represented as mean±SEM; n=8/group; *p<0.05 vs Young, #p<0.05 vs Old Control.

Ca^2+^ release from the sarcoplasmic reticulum (SR) via the cardiac ryanodine receptors (RyR2) triggers Ca^2+^-induced - Ca^2+^ release to induce cardiomyocyte contraction, this is followed by sequestration of Ca^2+^ via sarco/endoplasmic Ca^2+^-ATPase (SERCA2a) and Na^+^/Ca^2+^ exchanger (NCX) during cardiomyocyte relaxation^19^. Based on our observation of altered Ca^2+^ transients with aging and rapamycin treatment, we examine the expression and phosphorylation of these Ca^2+^ handling proteins^31^. The phosphorylation of S2808 of RyR2 is associated with Ca^2+^ leak and cardiac dysfunction^32,33^. We detected no differences in S2808 phosphorylation of RyR2 with aging or rapamycin treatment **(Supp. Figure 1A)**. Calsequestrin 2 (CSQ2) is a protein located at the junction of SR that functions as Ca^2+^ buffering system during Ca^2+^ release through RyR2. We detected no differences in the levels of CSQ2 in the three experimental groups **(Supp. Figure 1B)**. There were also no differences in the levels of SERCA2a amongst the three groups **(Supp. Figure 1C)**. Old hearts displayed increased expression of NCX compared to the young control hearts, and this age-related increase was normalized by rapamycin treatment **(Figure 2J)**.

### Rapamycin enhances myofibril relaxation kinetics and MyBP-C phosphorylation in old mice

In addition to changes in Ca^2+^ handling, alterations of myofilament properties modulate the contraction-relaxation kinetics of cardiomyocytes. To investigate if the improved cardiomyocyte relaxation kinetics of rapamycin treated old mice was associated with altered myofilament properties, we compared the properties of myofibrils from old control and old rapamycin treated mice following maximal (pCa 4.0) and sub-maximal (pCa 5.6) Ca^2+^ activation **(Table 1)**. There were no differences in the tensions generated by myofibrils from old control and old rapamycin treated mice at both conditions but rapamycin treatment increased the activation rate of force (*k*act) at sub-maximal Ca^2+^ activation, at Ca2+ levels seen during cardiomyocyte twitch contractions **(Table 1)**. The rates of fast phase of relaxation (*k*_REL, fast_) of myofibrils from old rapamycin treated mice were significantly faster compared to old controls at both pCa 4.0 and pCa 5.6 **(Table 1).** This may contribute to the improved cardiomyocyte relaxation kinetics in old rapamycin treated mice. The rate of the early linear, slow phase of relaxation (*k*_REL, slow_, correlated with cross bridge detatchment) and its duration (t_REL_,_slow_, correlated with thin filament deactivation) were not affected by rapamycin treatment **(Table 1)**.

**Table 1:**
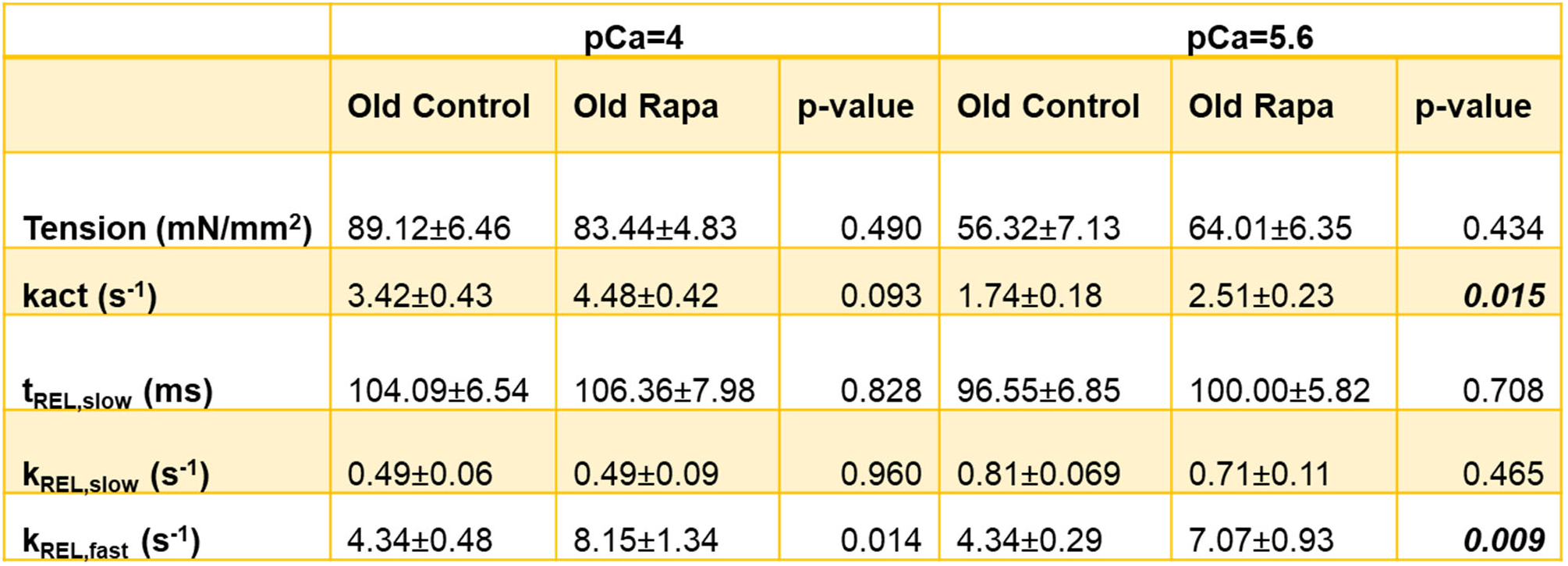
Tension generation and kinetic properties of old control and rapamycin-treated myofibrils following maximal (pCa 4.0) and submaximal (pCa 5.6) Ca^2+^ activation. Data represented as mean±SEM; n=10-11/group

|  | pCa=4 |  |  | pCa=5.6 |  |  |
| --- | --- | --- | --- | --- | --- | --- |
|  | Old Control | Old Rapa | p-value | Old Control | Old Rapa | p-value |
| Tension (mN/mm<sup>2</sup>) | 89.12±6.46 | 83.44±4.83 | 0.490 | 56.32±7.13 | 64.01±6.35 | 0.434 |
| k<sub>act</sub> (s<sup>-1</sup>) | 3.42±0.43 | 4.48±0.42 | 0.093 | 1.74±0.18 | 2.51±0.23 | **0.015** |
| t<sub>REL,slow</sub> (ms) | 104.09±6.54 | 106.36±7.98 | 0.828 | 96.55±6.85 | 100.00±5.82 | 0.708 |
| k<sub>REL,slow</sub> (s<sup>-1</sup>) | 0.49±0.06 | 0.49±0.09 | 0.960 | 0.81±0.069 | 0.71±0.11 | 0.465 |
| k<sub>REL,fast</sub> (s<sup>-1</sup>) | 4.34±0.48 | 8.15±1.34 | 0.014 | 4.34±0.29 | 7.07±0.93 | **0.009** |

Phosphorylation of cardiac troponin I (TnI), an inhibitory subunit of troponin, enhances the rate of cardiomyocyte relaxation^34^. Increased phosphorylation of TnI at Ser23/24 enhances cross bridge cycling rate and reduces Ca^2+^ sensitivity of steady-state force^35^. We observed reduced phosphorylation of TnI at S23/24 in old control and old rapamycin treated hearts compared to young counterparts, and rapamycin did not change the phosphorylation of Ser23/24 of TnI **(Figure 3A).** This suggests that reduced TnI phosphorylation at S23/24 may contribute to the age-related impairment in cardiomyocyte relaxation but not the rapamycin-induced improvement. Phosphorylation of myosin binding protein C (MyBP-C) promotes cross bridge cycling and diastolic function^36,37^. We observed that old control hearts displayed a trend towards a reduction in phosphorylation of MyBP-C at Ser282 site compared to young hearts **(Figure 3B)**. Most notably, rapamycin treatment substantially increased the phosphorylation of cMyBP-C at Ser282 in old rapamycin treated hearts **(Figure 3B)**, and this is consistent with the faster relaxation kinetics of myofibrils **(Table 1)** and cardiomyocytes of old rapamycin-treated mice **(Figure 1C)**.

**Figure 3:**
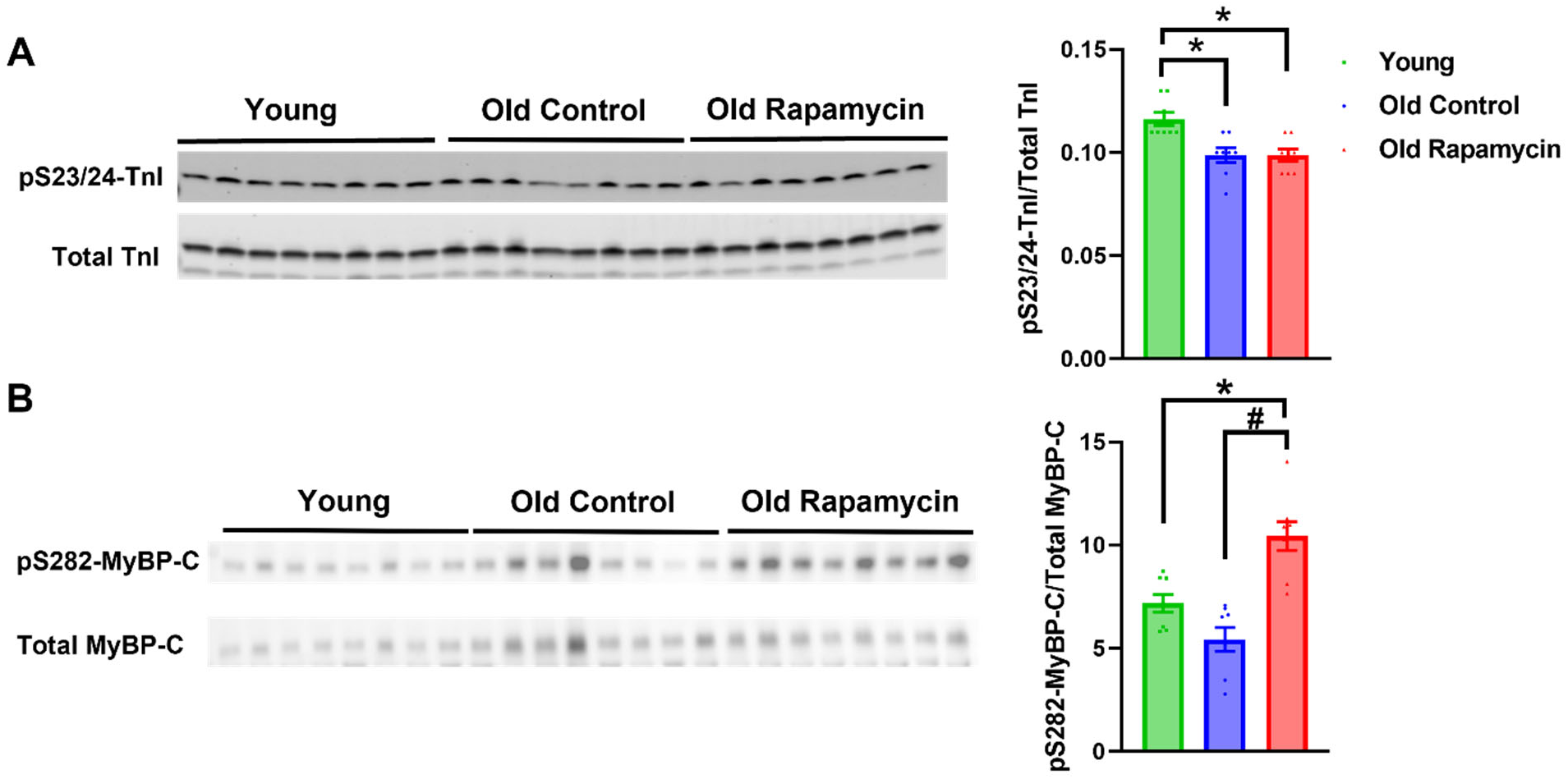
Aging and rapamycin treatment alter the phosphorylation of myofilament proteins. (A) Images of western blots of phosphorylated (S23/24) TnI and total TnI in heart tissues of young, old control and old rapamycin treated mice and the corresponding quantification. (B) Images of western blot of phosphorylated (S282) MyBP-C and total MyBP-C in heart tissues of young, old control and old rapamycin treated mice and the corresponding quantification. Data represented as mean±SEM; n=8/group; *p<0.05 vs Young, #p<0.05 vs Old Control.

### Rapamycin normalizes an age-related increase in myocardial stiffness via a mechanism independent of titin isoform shift

Passive stiffness of the myocardium is a key determinant of diastolic function and myocardial stiffness increases with age in mice, dogs and humans^38,39^. We measured the passive force-length relationship of demembranated trabeculae and detected significant increases in the passive force above 12% stretch in old trabeculae compared to young controls, indicating an age-related increase in myocardial stiffness **(Figure 4A)**. This age-related increase in myocardial stiffness was normalized by rapamycin treatment, as indicated by the reduced passive force in old rapamycin-treated mice compared to old control mice **(Figure 4A)**. Titin isoform ratios and titin phosphorylation regulate the passive stiffness of myocardium^20–24^. We measured titin isoforms in the heart to determine if titin isoform shift is responsible for the beneficial effect of rapamycin treatment. While there was an age-related increase in the N2BA/N2B ratio with age, rapamycin treatment did not alter titin isoform ratio **(Figure 4B)**. PKCα phosphorylates titin and increase passive myocardial stiffness^40,41^. Old rapamycin treated hearts exhibited reduced PKCα activation, represented by reduced PKCα-pS567 phosphorylation, compared to old control hearts **(Figure 4C)**. This suggests that rapamycin treatment may reduce passive stiffness by diminishing titin phosphorylation at the PKCα sites. Cardiac fibrosis also increases passive myocardial stiffness. Therefore, we examined whether rapamycin treatment altered fibrosis in old hearts. Trichome staining showed that old control and old rapamycin treated hearts had similar levels of collagen content **(Supp. Figure 2A-B)**. Myofilament Ca^2+^ sensitivity also modulates cardiomyocyte contraction-relaxation ^42^. We examined the force-Ca^2+^ relationship of demembranated trabeculae from young, old control and old rapamycin-treated mice to assess their Ca^2+^ sensitivities. We observed a small leftward shift in force-pCa curve **(Figure 4D)** and a trend towards a small increase in pCa_50_ in old controls compared to young control trabeculae **(Figure 4E)**. Trabeculae of old rapamycin treated mice exhibited a similar force-pCa curve **(Figure 4D)** and an unchanged pCa_50_ compared to old controls **(Figure 4E)**, indicating that rapamycin treatment did not affect the Ca^2+^ sensitivity of old myofilaments. The Hill coefficient, a measure of myosin cooperativity, did not change between the groups **(Supp. Table 1).**

**Figure 4:**
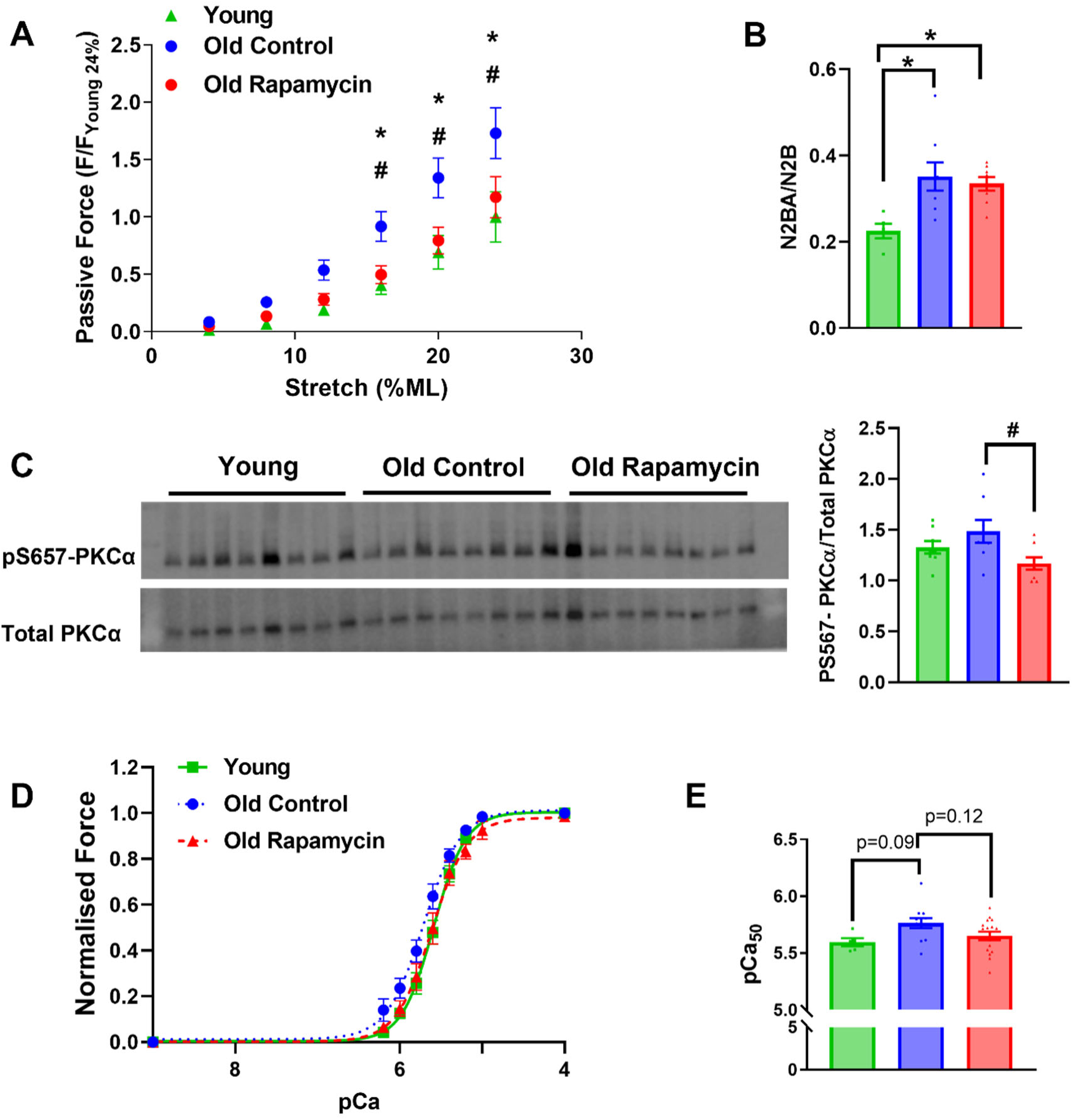
Rapamycin normalizes an age-related increase in myocardial passive stiffness via mechanisms independent of titin isoform shift. (A) Length-tension relationship of demembranated trabeculae from young, old control and old rapamycin treated mice. n=9-27/group. (B) The ratio of titin isoforms (N2BA/N2B) of the heart tissues of young, old control and old rapamycin treated mice. n=5-8/group. (C) Images of western blots of phosphorylated (S657) PKCα and total PKCα of heart tissues of young, old control and old rapamycin treated mice and the corresponding densitometry. n=8/group. (D) Force-pCa curves of demembranated passive trabeculae from young, old control and old rapamycin treated mice. (E) The pCa_50_ of demembranated trabeculae from young, old control and old rapamycin treated hearts. n=5-17/group. *p<0.05 vs Young, #p<0.05 vs Old control. All data represented as mean±SEM.

## Discussion

Age-related diastolic dysfunction is an unsolved medical problem with limited effective therapies^39,43^. Several structural and functional changes occur within the myocardium due to aging that have been linked to diastolic dysfunction^44^. Active relaxation of cardiomyocytes, a process that involves actin-myosin detachment and calcium reuptake, is a major determinant of diastolic function^45–47^. Passive stiffness of the myocardium is another key factor that regulates the diastolic reserve and function^47^. Rapamycin is a pharmacological inhibitor of mTORC1 and has been shown to reverse age-related diastolic dysfunction in mice^12,13^. However, the molecular mechanisms underlying the reversal of age-related diastolic dysfunction mediated by rapamycin treatment have remained unknown. In this study, we treated 24-month-old C57BL/6J mice with 10 weeks of rapamycin to investigate the mechanisms by which rapamycin reverses diastolic dysfunction with a focus on the changes in the contraction-relaxation machinery. Our work demonstrates that 1) rapamycin treatment normalizes the age-related impairments of sarcomere shortening and relaxation kinetics in cardiomyocytes; 2) age-related impairments in the kinetics of cardiomyocyte Ca^2+^ transients are partially restored by rapamycin treatment; 3) rapamycin treatment enhances the myofibril relaxation kinetics and promotes S282 phosphorylation of MyBPC in old mice; and 4) the reversal of age-related passive stiffness by rapamycin is independent of titin isoform shift.

### Age-related derangements in cardiomyocyte relaxation and Ca^2+^ handling is restored by rapamycin

We showed that old control cardiomyocytes exhibit lower sarcomere fractioning shortening with prolonged contraction (TPS_10_, TPS_50_ and TPS_90_) and relaxation times (RT_90_) compared to young cardiomyocytes **(Figure 1C, F and H-J)**. Importantly, these age-related derangements were partially or completely normalized by rapamycin treatment **(Figure 1C, F and H-J)**. The age-related impairment in cardiomyocyte relaxation and the normalization by rapamycin treatment are consistent with age-related diastolic dysfunction observed in old mice that can be reversed by rapamycin treatment^12,13^. Previous studies have shown that LV systolic function is relatively preserved or slightly reduced with aging at organ levels^13,48,49^. The age-related declines in sarcomere fractional shortening and contraction kinetics suggest that cardiac contraction is impaired at cardiomyocyte levels and that is partially normalized by rapamycin treatment.

Ca^2+^ cycling controls cardiac contraction and relaxation, and the remodeling of Ca^2+^ handling proteins is one of the major factors contributing to the mechanical and electrical dysfunction observed in HF^50^. Consistent with the prolonged cardiomyocyte contraction and relaxation times, old control cardiomyocytes displayed prolonged times to peak Ca^2+^ transient (TTP, TTP_50_, TTP_90_) and prolonged Ca^2+^ transient decay (DT_90_) compared to young cardiomyocytes **(Figure 2B and E-G)**. This observation is consistent with a previous study that showed slower time to peak Ca^2+^ transient and Ca^2+^ decay time in old cardiomyocytes compared to young cardiomyocytes^51^. In this study, we found that the age-related changes in DT_90_ and TTP_90_ are partially normalized by rapamycin treatment **(Figure 2B and F)**. These results suggest that rapamycin treatment attenuates the age-related derangements in Ca^2+^ handling to improve active cardiomyocyte contraction-relaxation. The complete normalization of age-related increase in RT_90_ **(Figure 1C)** and the partial normalization of age-related increase in DT_90_ **(Figure 2B)** by rapamycin treatment in old mice suggest that enhanced Ca^2+^ reuptake only partially contributes to the rapamycin-induced improvement in cardiomyocyte relaxation.

During diastole, cytosolic Ca^2+^ is removed by SERCA2-mediated Ca^2+^ reuptake in SR and NCX-mediated Ca^2+^ efflux out of the cell^19^. Previous studies have reported reduced or unchanged SERCA2 expression with age in murine hearts ^51–53^. In this study, we did not observe changes in SERCA2 expression with aging or rapamycin treatment. Reduced SERCA2 activity and increased oxidation and nitration of SERCA2 have been reported in aged mouse hearts^54–56^. Further studies are required to determine if rapamycin treatment reduces oxidation or nitration of SERCA2, decreases levels of SERCA2 inhibitor phospholamban, or enhances SERCA2 activity to improve cardiomyocyte Ca^2+^ transient and diastolic function. We detected increased NCX expression with aging, with rapamycin reversing this increase **(Figure 2J)**. NCX is known to be upregulated in failing hearts in both activity and expression^57,58^. This increase in NCX activity might be an adaptive mechanism to protect against diastolic Ca^2+^ overload^59^. We did not observe changes in RyR2 phosphorylation at S2808 and CSQ expression with aging or rapamycin treatment. Further investigations are needed to determine the mechanism by which rapamycin normalizes the slower TTP Ca^2+^ transient and contraction kinetics in old cardiomyocytes.

### Rapamycin enhances myofibril relaxation kinetics and MyBP-C phosphorylation in aging myocardium

There is ample evidence that HF degrades sarcomeric proteins that ultimately leads to sarcomere dysfunction^60–62^. Upon β-adrenergic stimulation, protein kinase A (PKA)-mediated TnI phosphorylation at S23/24 is associated with a decrease in myofilament Ca^2+^ sensitivity and contributes to an accelerated rate of cardiac relaxation^63^. Our data show a reduction in phosphorylation of TnI at S23/24 in old control hearts in comparison to young hearts and that the age-related decrease in S23/24 phosphorylation of TnI is not affected by rapamycin treatment (**Figure 3A**). MyBP-C is another sarcomeric protein that regulates contraction-relaxation by modulating cross bridge cycling. Mutations within MyBP-C are a common mechanism of hypertrophic cardiomyopathy and reduced MyBP-C phosphorylation has been observed in failing hearts. Using mouse models with transgenic expression of wild type or mutant MyBP-C proteins in MyBP-C null mice, Rosas et. al. showed a reduction in MyBP-C phosphorylation levels with aging in wild type MyBP-C expressing mice and that phosphorylation of MyBP-C mitigates age-related cardiac dysfunction by preserving myocardial relaxation ^64^. In this study, we show that aging has little to no effect on S282 phosphorylation levels of MyBP-C in the heart, but that rapamycin treatment significantly enhances S282 phosphorylation of MyBP-C (**Figure 3B**). The increase in S282 phosphorylation of MyBP-C in rapamycin treated old mice compared to old controls potentially contributes to the improved cardiomyocyte relaxation and diastolic function induced by rapamycin treatment. The difference in age-related changes in S282 phosphorylation of MyBP-C in our study compared to that observed by Rosas et al. may be due to different genetic backgrounds of mice used in the two studies.

### Rapamycin treatment rescues aging-associated increase in myocardial stiffness independent of titin isoform switch

We recently showed that rapamycin treatment reduces passive stiffness of the myocardium in aging mice^15^. In this study, we validated the age-related increase in myocardial stiffness and reversal with rapamycin **(Figure 4A)**, then examined the mechanism behind this effect. Titin is a sarcomeric protein that spans the Z disk to the M line of the sarcomere. Studies have shown that the ratio of titin isoforms, N2BA and N2B, determine titin’s contribution to myocardial stiffness, with reduced N2BA/N2B ratio resulting in increased stiffness^65,66^. We detected an isoform shift in the old control mice (increased N2BA/N2B) compared to young controls which is not affected by rapamycin **(Figure 4B)**. The unchanged N2BA/N2B ratio indicates that the reduced myocardial stiffness in old rapamycin-treated mice is mediated by mechanisms independent of titin isoform shift. Similar observation of increased passive stiffness despite increased N2BA/N2B ratio was observed in the heart of mice with diastolic dysfunction induced by transverse aortic constriction (TAC)^30^. The age-related increase in N2BA/N2B ratio is potentially a compensatory change to counteract the increased stiffness mediated by other mechanisms like titin phosphorylation and extracellular matrix (ECM) remodeling ^30^. Increased cardiac fibrosis and ECM deposition is a characteristic of cardiac aging and the age-related ECM remodeling contributes to the increased passive myocardial stiffness^67,68^. Rapamycin treatment did not alter collagen deposition measured by trichome staining **(Supp. Figure 2A-B)**. However, the effects of rapamycin on the expression of other ECM proteins remain to be determined.

In addition to titin isoform switching, phosphorylation of titin is another mechanism to regulate titin-based stiffness. PKCα phosphorylates titin in its PEVK region at 2 sites – S11878 and S12022 to increase tension^69^. Rapamycin treatment suppresses PKCα activation in the hearts of old mice, represented by S657 phosphorylation of PKCα^12^. In this study, we observed that rapamycin treatment significantly reduces the phosphorylation of S657 of PKCα in old hearts **(Figure 4C)**. This suggest that reduced PKC-mediated phosphorylation of titin may play a role in the reduced passive stiffness of old rapamycin treated hearts. This supports that aging and rapamycin may regulate passive myocardial stiffness by post-translational modification of titin.

The sarcomere function is regulated by both the thick (myosin) and thin (actin) filament regulatory pathways^70^. Cardiac performance is tightly regulated by sharp changes in force generation in dependence with myofilament Ca^2+^ sensitivity^71^. Myofilament activation is directly dependent on the amount of activating Ca^2+^, but is also determined by the myofilament Ca^2+^ sensitivity of force production^72^. An increase in myofilament Ca^2+^ sensitivity was observed in a canine model of heart failure ^73^, but no age-related differences in myofilament Ca^2+^ sensitivity were detected in aging rats^74^.In this study, we observed little to no change in Ca^2+^ sensitivity with age and unchanged Ca^2+^ sensitivity after rapamcyin treatment **(Figure 4D-E)**. This suggests that the regulation of myofilament Ca^2+^ sensitivity is not a mechanism by which rapamcyin improves diastolic function.

A limitation of this study is that only female mice were used. This is because the dose of rapamycin (14 ppm) used in this study has only been shown to reverse cardiac aging in female mice^12–14^. A higher dose (42 ppm) of rapamycin treatment has been shown to improve cardiac aging in male mice^15^. Future studies using the higher dose of rapamycin are required to investigate the mechanisms by which rapamycin improved cardiac function in old male mice.

### Conclusions and perspectives

Together, this study identified the age-related changes in the cellular and molecular modulators of diastolic function and revealed the impacts of late-life rapamycin treatment on these modulators. Importantly, we showed that rapamycin treatment enhances the cardiomyocyte relaxation kinetics, increases S282-MyBP-C phosphorylation, and reduces passive myocardial stiffness in the old murine heart. These mechanisms jointly contribute to the previously established rapamycin-induced reversal of age-related diastolic dysfunction. While rapamycin is only one of several new potential interventions to improve healthspan and longevity^75,76^, the mechanisms shown in this study may prove to be common elements of their beneficial effects on cardiac function.

## Acknowledgements

We acknowledge funding support from AHA 23POST1012408 to KAK, R35HL144998 to HG, K08HL128826 to FMH, P01AG001751 and P30AR074990 to MR, and K99/R00 AG051735 to YAC.

---

**Supplementary Figure 1.**
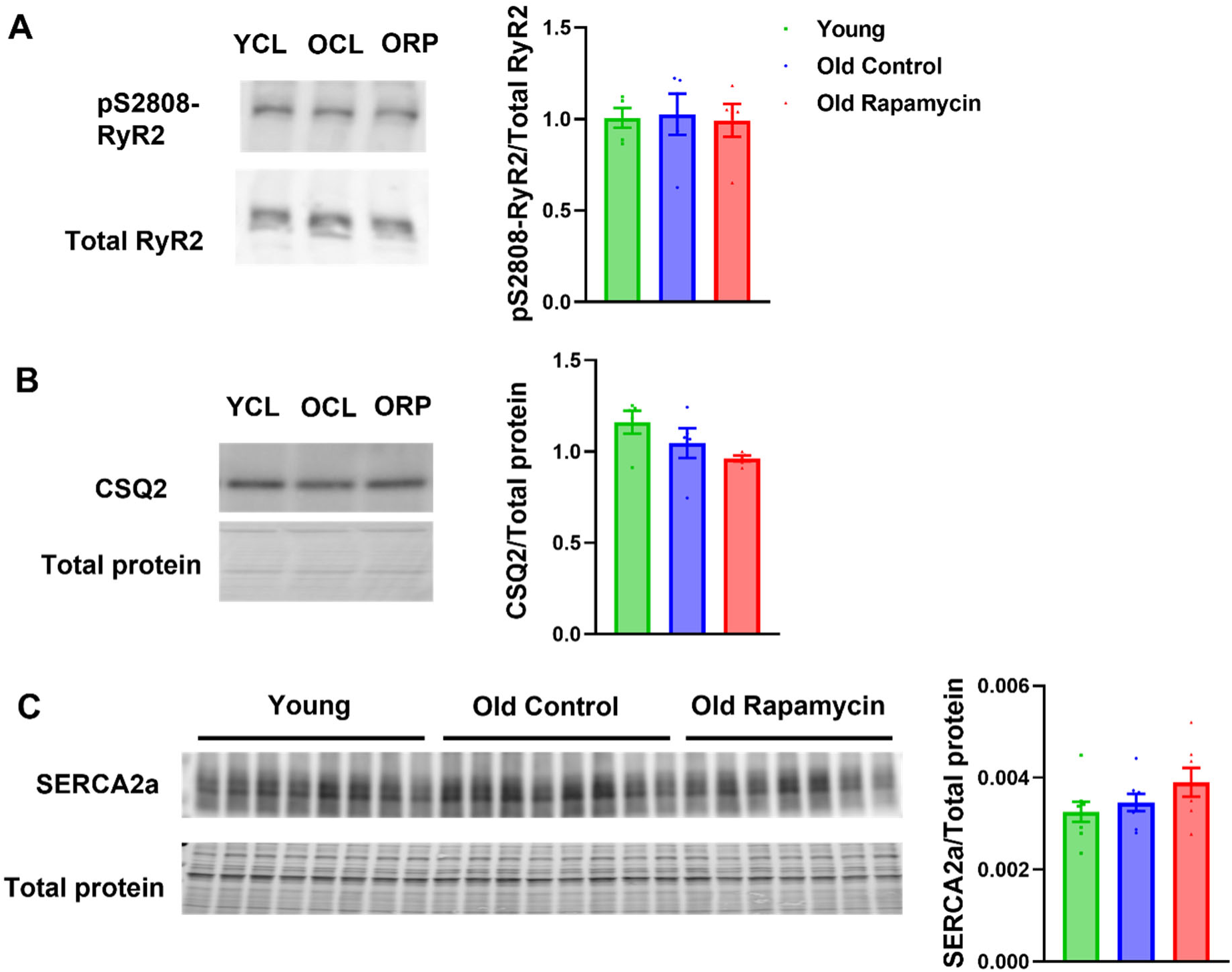
Aging and rapamycin treatment do not alter RyR2 phosphorylation and CSQ and SERCA2a expression. (A) Representative images of western blots of phosphorylated (S2808) RyR2 and total RyR2 of heart tissues of young, old control and old rapamycin treated mice and the corresponding quantification. n=5/group. (B) Representative images of CSQ2 western blot and total protein staining of heart tissues of young, old control and old rapamycin treated mice and the corresponding quantification. n=5/group. (C) Images of SERCA2a western blot and total protein staining of heart tissues of young, old control and old rapamycin treated mice and the corresponding quantification. n=7-8/group. All data represented as mean±SEM.

**Supplementary Figure 2.**
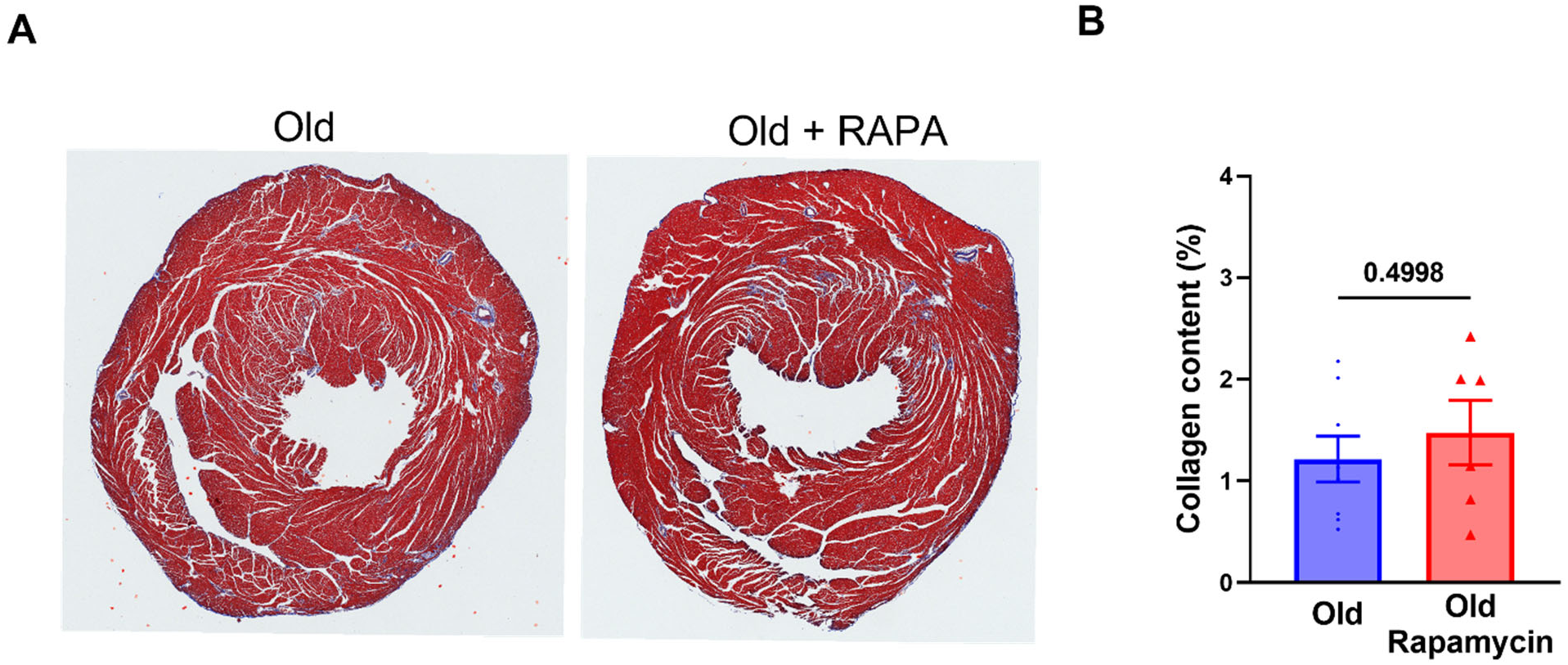
Rapamycin treatment do not alter collagen content. (A) Representative images of trichome staining of heart tissues of old control and old rapamycin treated mice and (B) the corresponding quantification. Data represented as mean±SEM; n=7-8/group.

**Supplementary Table 1.** Summary table of force-pCa measurements of demembranated trabeculae from young, old control and old rapamycin treated mice. Data represented as mean±SEM; n=5-17/group.

|  | Young (n=5) | Old Control (n=12) | Old Rapa (n=17) |
| --- | --- | --- | --- |
| **F<sub>max</sub>** (mN/mm<sup>2</sup>) | **28.47±5.35** | **45.24±5.96** | **41.00±5.87** |
| **pCa<sub>50</sub>** | **5.59±0.04** | **5.76±0.04** | **5.65±0.04** |
| **h(Hill Coefficient)** | **2.33±0.12** | **2.20±0.14** | **2.44±0.14** |

